# ppGpp functions as an alarmone in metazoa

**DOI:** 10.1101/2019.12.16.878942

**Authors:** Doshun Ito, Hinata Kawamura, Akira Oikawa, Yuta Ihara, Toshio Shibata, Nobuhiro Nakamura, Tsunaki Asano, Shun-Ichiro Kawabata, Takashi Suzuki, Shinji Masuda

## Abstract

Guanosine 3’,5’-bis(pyrophosphate) (ppGpp) functions as a second messenger in bacteria to adjust their physiology in response to environmental changes. In recent years, the ppGpp-specific hydrolase, metazoan SpoT homolog-1 (Mesh1), was shown to have important roles for growth under nutrient deficiency in *Drosophila melanogaster*. Curiously, however, ppGpp has never been detected in animal cells, and therefore the physiological relevance of this molecule, if any, in metazoans has not been established. Here, we report the detection of ppGpp in *Drosophila* and human cells and demonstrate that ppGpp accumulation induces metabolic changes, cell death, and eventually lethality in *Drosophila*. Our results provide the first evidence of the existence and function of the ppGpp-dependent stringent response in animals.

## Main Text

Organisms must adjust their physiology in response to environmental changes. The stringent response is one of the most important starvation/metabolic control responses in bacteria and is controlled by the hyper-phosphorylated nucleotide, ppGpp (*1*, *2*). ppGpp accumulates in bacterial cells upon exposure to various stresses, and the accumulated ppGpp functions as an alarmone that can alter transcription (*3*–*6*), translation (*7*, *8*) and certain enzymatic activities (*6*, *9, 10*) to overcome a stress (*3*). In *Escherichia coli*, ppGpp level is regulated by two distinct enzymes, namely RelA and SpoT (11). Both RelA and SpoT catalyze pyrophosphorylation of GDP (or GTP) by using ATP to produce ppGpp (*12*), and SpoT, but not RelA, hydrolyzes ppGpp (*13*). The RelA/SpoT homologs (RSHs) are universally conserved in bacteria (*14*) and have pivotal roles in various aspects of bacterial physiology, including the starvation response (*1*, *2*), growth-rate control (*15*), antibiotic tolerance (*16*), and darkness response (*17*). RSHs are also found in eukaryotes, including the green algae *Chlamydomonas reinhardtii* (*18*) and land plant *Arabidopsis thaliana* (*19*) as well as metazoa, including humans and *Drosophila melanogaster* (*14*, *20*). Metazoan SpoT homolog-1 (Mesh1) is one class of RSHs found in metazoa. Although most RSHs have both ppGpp synthase and hydrolase domains, Mesh1 has only a ppGpp hydrolase domain (Fig. 1A). Mesh1 hydrolyzes ppGpp in vitro with comparable efficiency to bacterial RSHs (*21*, *22*). The *Drosophila Mesh1 loss-of-function mutant* (*Mesh1 lof*) exhibits retarded growth, especially during starvation (*21*), suggesting that the ppGpp-dependent stringent response has been conserved in metazoa. However, this hypothesis is still under investigation because no known-ppGpp synthase has been identified and ppGpp itself has never been detected in animals (*21*).

**Fig. 1.**
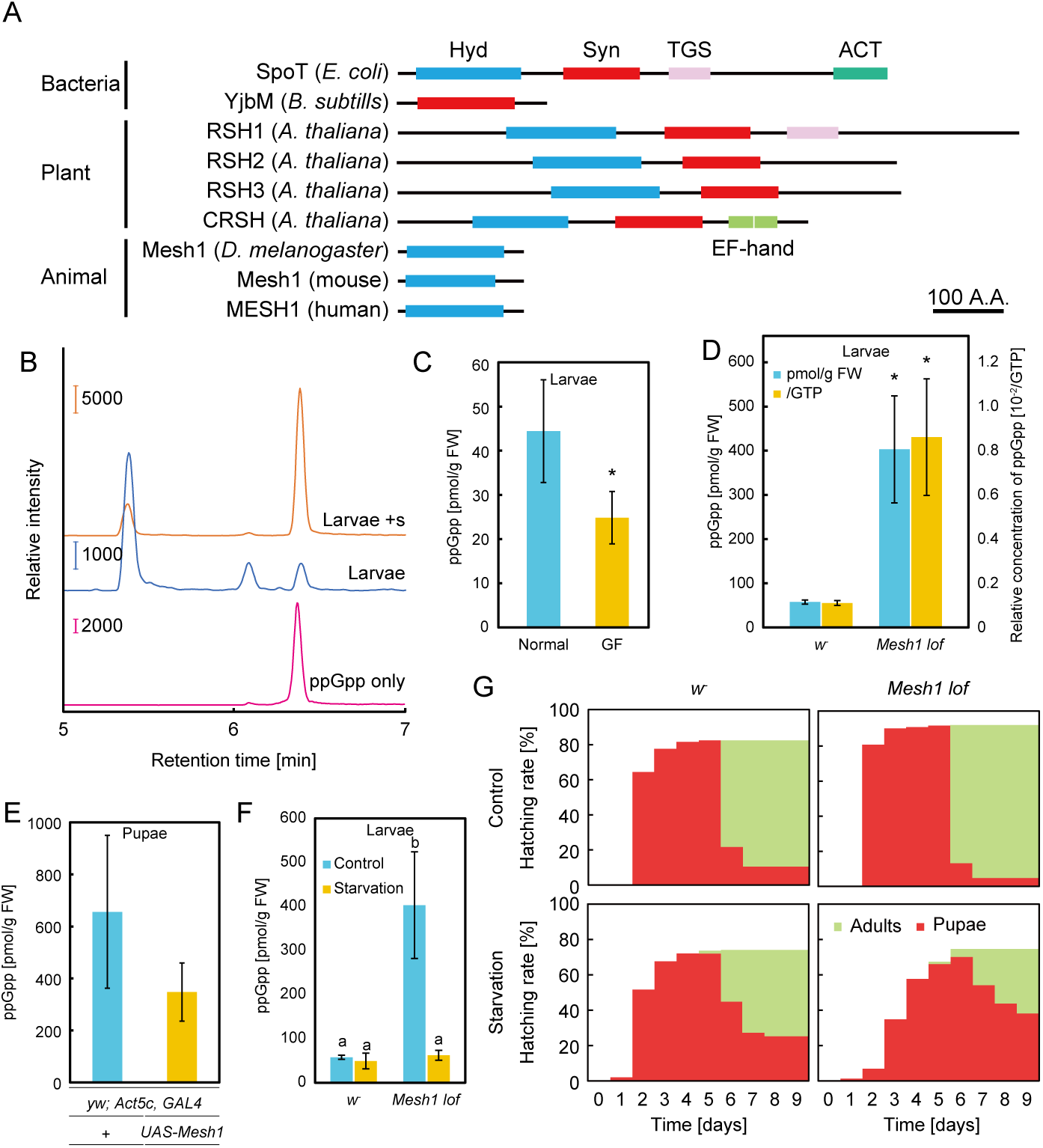
ppGpp detection in *Drosophila*, and *Mesh1* mutant phenotypes. **(A)** Schematics of the primary structures of RSHs. Hyd: (p)ppGpp hydrolase domain; Syn: (p)ppGpp synthase domain; ACT: aspartate kinase chorismite mutase TyrA domain; EF hand: Ca^2+^ binding helix E and F motif; TGS: threonyl-tRNA synthetase, GTPase, SpoT domain; A.A.: amino-acids. **(B)** Multiple-reaction-monitoring chromatograms for guanosine tetraphosphate ions in extracts from *w*^−^ third instar larvae. The symbol + denotes the exogenous addition of the ppGpp standard to the sample. Concentrations of ppGpp in *w*^−^ instar larvae grown under normal and germ-free (GF) conditions **(C)**, in *Mesh1 lof* late third instar larvae **(D)**, in *Mesh1 gof* day 2 pupae **(E)**, and in *Mesh1 lof* under normal and starvation conditions **(F)**. Values represent the mean ± S.D. (*n* =3). \**p* < 0.05, Student’s t-test. Different letters indicate significant differences (*p* < 0.05; Tukey’s test). **(G)** Hatching rate of the larvae (4 days after egg laying) of *w*^−^ and *Mesh1 lof*, which were initially grown on standard medium and then transferred to normal (Control) or starvation conditions. The colors indicate the number of flies from each stage (%); red: pupae, green: adults.

To check for the existence of ppGpp in animals, we applied the liquid chromatography-tandem mass spectrometry (MS)–based ppGpp quantification method, which we previously established for characterization of ppGpp function in plants (*23*). We extracted nucleotide pools from *Drosophila* late third instar larvae, in which *Mesh1* expression was higher than at other stages as examined by northern blotting (*21*). Using the guanosine tetraphosphate–specific MS/MS mode (*m/z* transition from 602 to 159), three peaks were observed at 5.4, 6.1 and 6.4 min (Fig. 1B, blue), similar to what was observed for the detection of ppGpp in *Arabidopsis* tissues (*23*). To determine which peak, if any, contained ppGpp in the *Drosophila* extract, we added ppGpp to the larvae extract (final conc. 16.7 nM) and subjected the sample to MS/MS analysis. The elution profile of the mixed sample revealed an increased area of the 6.4-min peak (Fig. 1B, orange) that matched the ppGpp standard peak (Fig. 1B, pink), indicating that it corresponded to ppGpp elution, as also seen in *Arabidopsis* (*23*). The 5.4-min peak was assigned as GTP (owing to cross-talk of the product ion), and the 6.1-min peak represented an unknown molecule (Fig. S1). Given the unknown molecule was detected in the ppGpp and GTP standards (Fig. 1B, pink; Fig. S1), it was contaminated in the products.

Using this MS/MS method, we quantified ppGpp concentrations in various developmental stages of wild-type (WT) *Drosophila* (*w*^−^): eggs from the strain Canton-S (CS), late third instar larvae (male and female), pupae (day 2), virgins (male and female), and day 4 adults (male and female). ppGpp was detected in extracts from all stages (Fig. 1B, S2A-D) at 50–250 pmol/g fresh weight (FW) (Table S1), which is comparable to that found in *Arabidopsis* (~100 pmol/g FW) (*23*). Interestingly, the amount of ppGpp in pupae was greater than that at other stages (Fig. S3A, B, Table S1). Given that *Mesh1* is highly expressed in larvae and suppressed in pupae (*21*), ppGpp tends to be degraded in the larval stage, and low expression of *Mesh1* in the pupal stage induces ppGpp accumulation. The concentration of ppGpp (per FW) in female adults was slightly, but significantly, higher than in male adults (Fig. S3A); however, the relative proportion of ppGpp to GTP in females was ~50% less than that observed in males (Fig. S3B). The GTP concentration in mating females was also higher (~3-fold) than in mating males (Fig. S3C), although this concentration difference was not found in adult virgins (Fig. S3A, B), suggesting that the increased ppGpp and GTP in mating females was derived from fertile eggs.

Bacterial cells contain a large amount of ppGpp (~470 nmol/g FW) (*24*, *25*), and animals harbor symbiotic bacteria (*26*), suggesting that the ppGpp detected in *Drosophila* may have originated from such bacteria. To investigate this hypothesis, we quantified ppGpp in germ-free *Drosophila*, which lack bacterial symbionts (*27*). Indeed, ppGpp was clearly detected in germ-free *Drosophila*, although the concentration was significantly lower than in non-germ-free control flies (Fig. 1C, S4). These results indicated that, although ~50% of the detected ppGpp might have originated from symbiotic bacteria (Fig. 1C), some ppGpp is synthesized by an unknown enzyme(s). To confirm this, we detected ppGpp in extracts of germ-free human cells (HeLa and PEAKrapid) and *Drosophila* (S2) cells in culture. The chromatograms showed peaks at the same retention times as those of late third instar larvae of *Drosophila* (Fig. S4). The ppGpp concentration in HeLa cells was ~40 pmol/g FW, and the relative concentration of ppGpp to GTP in both HeLa and S2 cells was ~2.5 × 10^−3^ (Table S1), which is comparable to that determined in *Drosophila* larvae.

Mesh1 expressed in *E. coli* exhibits ppGpp hydrolase activity (*21*, *22*). However, Mesh1 activity *in vivo* has never been measured because ppGpp has not been detected in metazoa (*21*). To verify that metazoan Mesh1 has ppGpp hydrolase activity, we quantified ppGpp in extracts of *Drosophila Mesh1 lof* and the *Mesh1*-overexpressing transformant (*Mesh1 gof*: gain-of-function) (*21*). The chromatograms of ppGpp extracted from *Mesh1 lof* had a sharp ppGpp peak (Fig. S2E, F), and the concentration of ppGpp in larvae was significantly (~7-fold) higher than in WT (Fig. 1D, Table S1). The *Mesh1 lof* pupae contained ~3-fold more ppGpp than did WT (Fig. S3D). To induce *Mesh1* expression in *Drosophila*, we applied a GAL4/UAS system (*28*). We crossed *UAS-Mesh1* transformants with flies expressing GAL4 and used an actin promoter, *Act5c*, for systemic and constitutive overexpression of *Mesh1* (*Mesh1 gof*). The amount of ppGpp in *Mesh1 gof* was ~350 pmol/g FW in pupae, which was ~50% that in the control (*P* = 0.21, *t*-test) (Fig. 1E). These results strongly suggested that Mesh1 has ppGpp hydrolase activity *in vivo*. We also found that GTP levels in *Mesh1 lof* larvae and pupae were approximately 10% and 20% lower, respectively, than those in WT (*w*^−^) (Fig. S3E), suggesting that ppGpp plays a role in GTP homeostasis, although GTP levels in *Mesh1 gof* pupae were the same as those in WT (Fig. S3F).

To further investigate ppGpp function in *Drosophila*, we characterized the *Mesh1 lof* mutant phenotype during hatching from larvae to pupae and from pupae to adults under starvation conditions. On rich medium (control), most larvae and pupae hatched at the second and sixth day, respectively, after transfer to the medium, and final hatching rates from larvae to adults were 72% for *w*^−^ and 87% for *Mesh1 lof* (Fig. 1G). Interestingly, on the starvation medium, hatching of *Mesh1 lof* was delayed compared with *w*^−^, and the hatching rate from larvae to adults in *Mesh1 lof* was only 37% at the ninth day, whereas in *w*^−^ it was 49% (Fig. 1G). These results indicated that Mesh1-dependent control of ppGpp concentration is important for growth and the response to starvation. We also measured the amount of ppGpp in *Drosophila* after transfer to starvation conditions. We used late third instar larvae (shown in Fig. 1D) as the control, and larvae at day 4 after-egg-laying were grown in rich medium and then transferred to the starvation medium and incubated for one day for comparison with late third instar larvae. The starved *w*^−^ larvae weighed significantly less than the *Mesh1 lof* larvae (Fig. S5A). The ppGpp concentration in starved *w*^−^ larvae did not differ from that of the control, whereas that in *Mesh1 lof* larvae was lower, i.e., the same as the control upon starvation (Fig. 1F, Fig. S5B, C). The GTP level (per FW) in *Mesh1 lof* remained lower than in *w*^−^, although the GTP level in *w*^−^ increased upon starvation (Fig. S5D). These results suggested that the lower GTP level in *Mesh1 lof* was a consequence of an increase in ppGpp, and the lower GTP level under normal conditions induced the pupation delay (Fig. 1G, Fig. S3E).

To further investigate the function of ppGpp in *Drosophila*, we heterologously expressed *yjbM* of *Bacillus subtilis* that encodes a small ppGpp synthase (Fig. 1A) (*29*). We produced a *UAS-YjbM* line, which drives *yjbM* expression constitutively when it is crossed with the actin-specific driver-line *Act5c-GAL4*. We found that offspring having both *UAS-yjbM* and *Act5c-GAL4* constructs were lethal, and progeny could not be produced (data not shown). As an alternative, we transiently expressed *yjbM* under the control of the heat-shock-driven FLPase system (*hs-yjbM*) at day 3 after-egg-laying, when larvae show strong resistance against environmental changes (*30*). *hs-yjbM* larvae could survive for several days after the induction of *yjbM*. The ppGpp chromatogram of the *hs-yjbM* exhibited a strong peak at 6.4 min, which corresponds to the ppGpp retention time of the control sample (Fig. 2A), and indicated ~1,200-fold higher ppGpp than that in the control (Fig. 2B). All the *hs-yjbM* died during the larvae and pupae stages (Fig. 2C), indicating that hyper-accumulation of ppGpp induces lethality in *Drosophila*, similar to that observed in the *E. coli spoT* mutant (*31*).

**Fig. 2.**
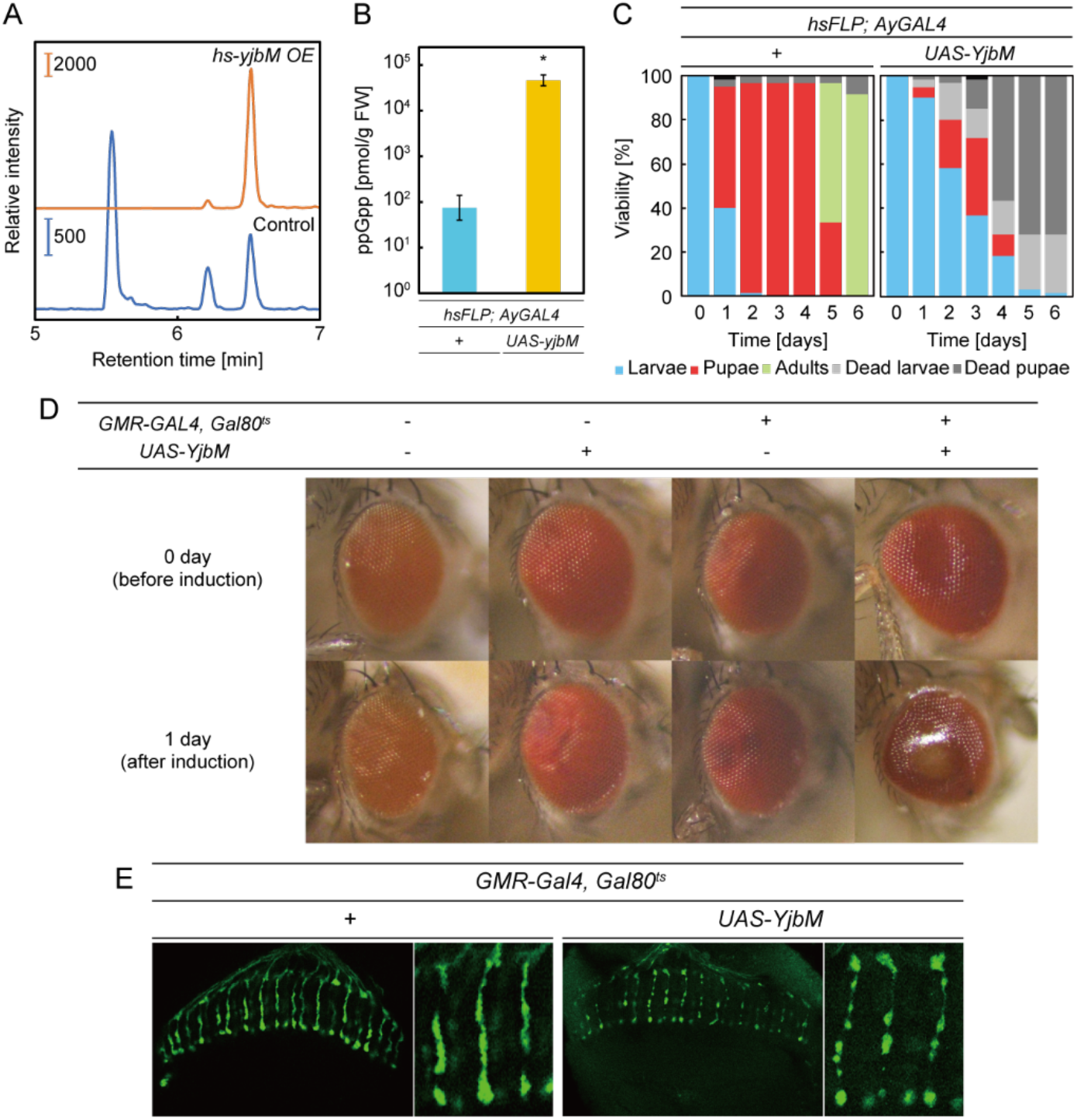
Excess ppGpp inhibits growth and promotes cell death. **(A)** Guanosine tetraphosphate– specific MS chromatograms of nucleotides extracted from *hs-yjbM OE* (a heat-inducible *yjbM* overexpression line) and control larvae. **(B)** The concentration of ppGpp extracted from *hs-yjbM OE* larvae and the control (1 day after induction). Values represent the mean ± S.D. (*n* =3). \**p* < 0.05, Student’s t-test. **(C)** *Drosophila* development after induction of ppGpp production. Values represent the mean (*n* =3). The colors indicate each stage of flies; blue: larvae, red: pupae, green: adults, light gray: dead larvae, gray: dead pupae. **(D)** Optical images of eyes and **(E)** confocal fluorescence images of axons (visualized with red fluorescent protein) of flies overexpressing *yjbM* by the eye-specific *GMR* promoter. Flies incubated at 18 °C one day after emergence (before induction; upper) were transferred to 31 °C to induce overexpression of *yjbM* in the eye. Images were acquired one day after induction.

Because systemic expression of *yjbM* is lethal in *Drosophila*, we could not further investigate the influence of ppGpp accumulation. Thus, we conducted site-specific expression of *yjbM* by crossing with the eye-specific driver-line *GMR-GAL4* (*GMR-YjbM*). We also used the temperature-dependent GAL4 repressor GAL80^ts^ to reduce leaky expression of YjbM. Before induction of *yjbM* expression, the surface of the eyes was smooth, but abnormally glossy (Fig. 2D), suggesting partial induction of cell death perhaps owing to leaky expression of *yjbM*. After induction of *yjbM*, the centers of the eyes became more yellow, supporting the cell death hypothesis. Microscopic analysis of the optic nerves inside the eyes revealed fragmentation of the axons (Fig. 2E). Similar axon fragmentation has been reported in axon degeneration and neural cell death (*32*), suggesting that excess ppGpp is cytotoxic. On the other hand, the eye morphology of *Mesh1 lof* did not differ from that of *w*^−^, as seen in both optical images and confocal images of dissected eyes (Fig. S6).

We next performed metabolome analysis of *Mesh1 lof* and *Mesh1 gof* to investigate the effects of ppGpp reduction and/or accumulation on metabolism. The levels of certain metabolites were significantly altered in the mutants compared with controls (Fig. 3, Fig. S7, Table S2, S3). In *Mesh1 lof*, TCA cycle metabolites including 4-hydroxybenzoate, fumarate and malate were higher than in WT (CS). In *Mesh1 lof*, the level of Arg was lower and that of GABA was higher than in CS (Table S2). In the transsulfuration pathway of *Mesh1 lof*, the CySSG level was higher and taurine level lower than in the control (Fig. 3, Table S2). In *Mesh1 gof*, the two arginine metabolites citrulline and creatine were present at lower levels than in the control (Fig. S7A, Table S3). In the methionine cycle and the transsulfuration pathway, methionine and cystathionine levels were lower, whereas adenosine and pyroglutamic acid levels were higher (Fig. S7A, Table S3). The levels of only six metabolites were altered in each of *Mesh1 lof* and *Mesh1 gof*, and three of them, including the two TCA cycle metabolites malate and fumarate, were higher in *Mesh1 lof* and lower in *Mesh1 gof* (Fig. 3, Fig. S7, Table S2, S3). These results imply that cellular ppGpp concentration influences both the TCA cycle and urea cycle in mitochondria. In *Mesh1 gof*, carboxymethyl lysine and pyroglutamic acid, one of the advanced glycoxidation end products and a glutathione metabolite, respectively, were increased (Fig. S7A, Table S3). Advanced glycoxidation end products are involved in aging and in the development of many degenerative diseases in humans (*33*), suggesting that the basal level of ppGpp is important for maintaining metabolic homeostasis in animal cells.

**Fig. 3.**
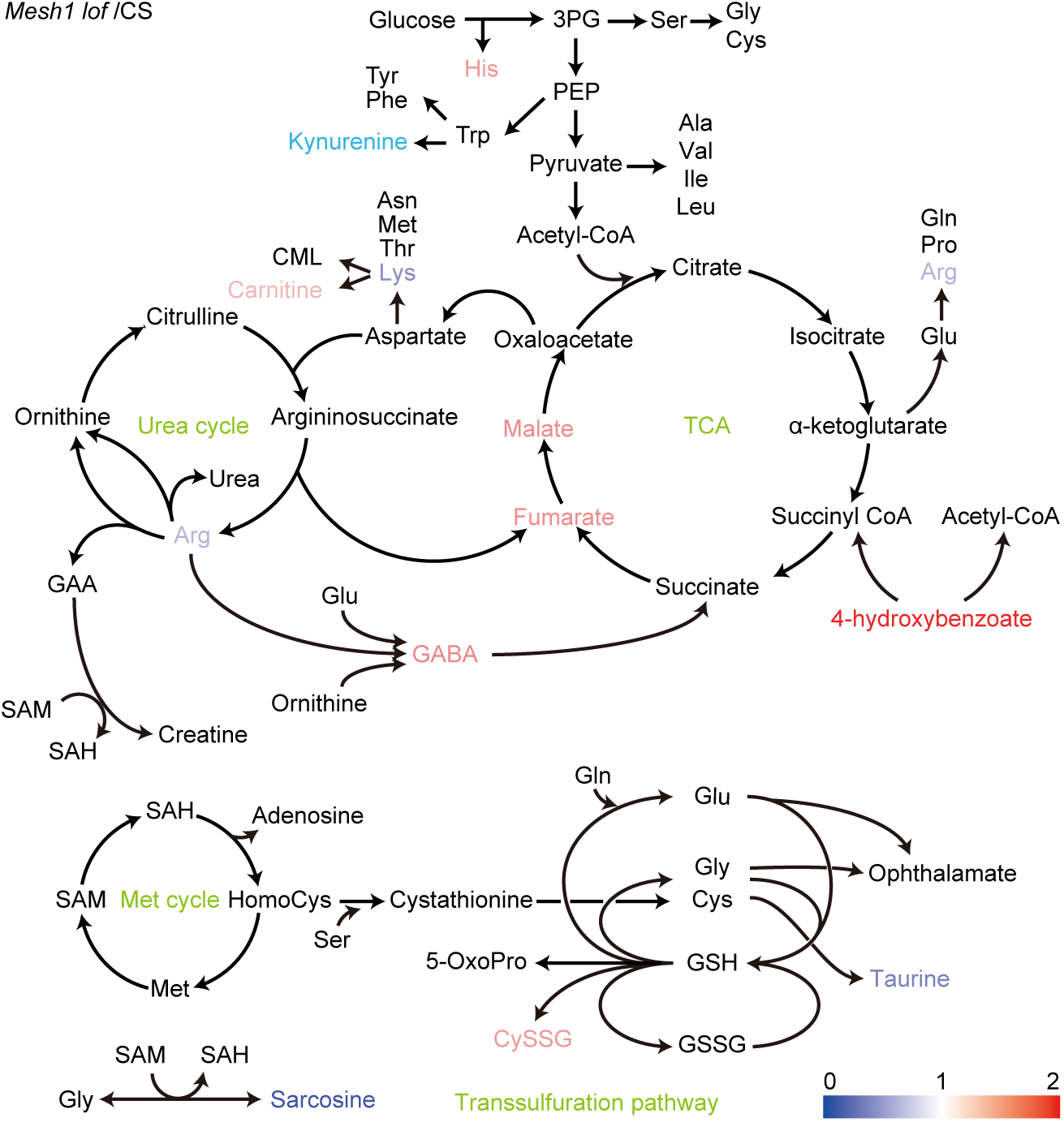
Statistically significant changes in the concentrations of metabolites in *mesh1 lof* larvae compared with concentrations in *w*^−^ larvae (*p* < 0.05; *n* = 6). Fold-change values in metabolite accumulation are expressed along a color gradient. 3PG: 3-phosphoglyceric acid, 5-OxoPro: pyroglutamic acid, CML: carboxymethyl lysine, CysSG: cysteine-glutathione disulfide, GAA: guanidinoacetic acid, GABA: gamma-aminobutyric acid, GSH: glutathione, GSSG: glutathione disulfide, SAM: S-adenosylmethionine, SAH: S-adenosylhomocysteine, and PEP: phosphoenolpyruvate.

It is well established that ppGpp is a bacterial and plastidial second messenger. Our results reveal that metazoa also use the molecule to modulate metabolism during starvation. Although the metazoan ppGpp synthase as well as exact ppGpp targets have not been identified, our results suggest that enzymes involved in purine metabolism (*34*) are directly regulated by ppGpp. In *Mesh1 gof,* adenosine and adenine levels were higher and hypoxanthine level was lower than those in the control (Table S3). In *B. subtilis*, (p)ppGpp directly inhibits certain enzymes involved in GTP synthesis, including guanylate kinase and hypoxanthine phosphoribosyltransferase (3), and this inhibition decreases the cellular GTP pool and retards transcription by reducing the concentrations of factors (e.g., ribonucleotides) necessary for transcription (*35*). The reduced levels of hypoxanthine observed in *Mesh1 gof* (Table S3) were likely due to the reduction of ppGpp (Fig. 1E), and metazoan hypoxanthine phosphoribosyltransferase conserves the amino-acid residues necessary for ppGpp binding in bacterial enzymes (*36*). Together with the increased concentration of GTP in the *Mesh1 lof* (Fig. S3E), these results imply that ppGpp in metazoans regulates the GTP/ATP ratio by controlling purine-nucleotide metabolism, as in bacteria. Moreover, other GTP-binding proteins could be ppGpp targets (*37*). Overall, our new method for quantifying ppGpp in animal cells should provide new opportunities for investigating the stringent response in animals.

## Acknowledgments

We thank J. Chung (Seoul National University) for proving fly strains.

## Author contributions

D.I. and S.M. designed and supervised the project. D.I. and Y.I. quantified ppGpp and GTP. D.I., H.K. and T.S. constructed and characterized *Drosophila* mutants. A.O. performed metabolome analysis. T.S. and S-I.K. grew and cultivated germ-free flies.

N.N. and T.A. maintained and sampled cultured cells. D.I. and S.M. wrote the first draft of the manuscript. All authors contributed to the discussion of the results.

## Data and materials availability

All data are presented in the manuscript or the supplementary materials.

## Supplementary Materials

Materials and Methods

Figures S1-S6

Tables S1-S3

