## Supplemental Figures and Tables for "ppGpp functions as an alarmone in metazoa"

**This PDF file includes:**

Materials and Methods
Figs. S1 to S7
Tables S1 to S3

### Materials and Methods

#### Fly strains, genetics, and growth conditions

Flies were kept in standard *Drosophila* medium at 25 °C, except for activating $GAL80^{ts}$ (18 °C). To activate the heat-shock (*hs*) promoter, flies were transferred from 18 °C to 31 °C after one day from emergence. To calculate ppGpp concentration, each stage of *Drosophila* was collected as follows; eggs lied on an apple juice medium for 19 h, the third instar larvae, pupae at 2 days after pupation, virgins, and adults at 4 days after hatching. To induce *yjbM* overexpression on day 3 after egg lying (AEL), *hs-yjbM* was heat-shocked for 30 min at 37 °C and then held at 10 min at room temperature, and this heat-shock procedure was repeated. To compare the pupation rate and ppGpp level under starvation conditions, eggs from flies of each genotype were collected for 2 h on an apple juice medium. Eggs were aged on standard *Drosophila* medium at room temperature (25 °C), and the standard medium was used as the normal growth condition (see below for recipe). To assess the effects of starvation, larvae at 4 days AEL under medium were transferred to starvation medium (see below for recipe). The following fly stocks and mutant alleles were used: CS, $w^{1118}$ ($w^{-}$), *Mesh1 loss-of-function* mutant (*Mesh1 lof*), *UAS-Mesh1*, *UAS-yjbM*, *Act5C-Gal4*, *Act5c-Gal4-tubGAL80*$^{ts}$, *hsFLP-AyGAL4-UAS-GFP*, and *GMR-myr-RFP-GMR-GAL4-tubGal80*$^{ts}$. The *UAS-yjbM* (residues 1–636 with a C-terminal FLAG tag) fly was produced with the *Φ*C31 system and plasmid pUASTattB (*38*), which was integrated into the specific site (VK31) of chromosome III. Transformants were produced by BestGene Inc. (U.S.A). *Mesh1 lof* and *UAS-Mesh1* were kindly provided by J. Chung (Seoul National University) (*21*). CS (Canton-S) was used for detection of ppGpp from eggs as the wild-type strain. $w^{-}$ was used as the control strain for comparison with mutants. Eye-specific expression was achieved using the GAL4-UAS expression system (*28*). *GMR-GAL4* was used to induce expression in eyes (*39*). To analyze the *yjbM* overexpression (OE) line, we made +/+; +/+; *Act5c-GAL4, Tub-Gal80$^{ts}$/UAS-yjbM* (or +) (*yjbM OE*), *hs-FLP/w; AyGAL4, UAS-GFP/+; /UAS-yjbM* (or +) (*hs-yjbM OE*), and *GMR-myr-RFP/w; GMR-GAL4, tub-Gal80$^{ts}$/+; +/UAS-yjbM* (or +) (*GMR-yjbM OE*). To analyze the *Mesh1 gof*, we crossed *UAS-Mesh1* flies with *yw; Act5c-Gal4/TM6B*.

Gene expression

To evaluate the effect of ppGpp accumulation, we used the GAL4/UAS system (*28*). We used transformants that had *UAS-yjbM* or *UAS-Mesh1*, which were silent in the absence of GAL4. They were crossed with flies expressing GAL4. In the progeny of this cross, GAL4 protein activated transcription of *yjbM* or *Mesh1* by binding to an UAS (*28*).

*Glass multimer reporter (GMR)-GAL4* was used for induction of photoreceptor gene expression. The GMR sequence is an engineered transcription regulatory region containing a binding site for the protein gl, a photoreceptor-specific transcription factor (*39*). *myr* is the sequence from *D. melanogaster Src64B* that includes the myristoylation signal and is used to target a protein to the plasma membrane.

To induce *yjbM* expression at a specific developmental time, we used *GAL80$^{ts}$* or the flippase (flp)/FRT system in conjunction with the GAL4/UAS system, which was controlled by the *hs* promoter (*30*, *40*). *GAL80$^{ts}$* is temperature-sensitive GAL80$^{ts}$ gene. At 18 °C, GAL80$^{ts}$ binds GAL4, and this heterodimer works as a GAL4 repressor. Upon transfer to a temperature above 29 °C, GAL80$^{ts}$ is unable to bind GAL4, and its repressive function is lost (*40*). Thus, from routine culture at 18 °C, flies were transferred to 31 °C when induction of *yjbM* was desired. With the flp/FRT system, the transformants had three genes to control gene expression: heat shock-*flp* fusion gene, the Act5C promoter-GAL4 fusion gene with a FRT cassette containing a transcription termination signal, and *UAS-yjbM*. Before heat shock, *Act5C-GAL4* was interrupted by the transcription termination signal in the FRT cassette. Heat shock induced expression of *flp* encoding the flippase, which excised the FRT cassette from the fusion gene. The GAL4 protein expressed from the rearranged fusion gene activated transcription of *UAS-yjbM* (*30*).

Culture medium for *Drosophila*

For the standard medium, 15 g soybean flour, 10 g agar, 100 g cornmeal, 30 g yeast, 30 g malt, and 50 g glucose were mixed in 1 l water and then heated at 90 °C for 1.5 h. After the mixture had cooled, 2 ml propionic acid and 2.5 g nipagin in 5 ml 100% ethanol were added. The mixture was then dispensed into columned vials or bottles until it covered the bottom to a depth of ~2 cm. The starvation medium consisted of 20% (w/v) sucrose-agar medium containing 0.5% ethanol and 0.2% propionic acid in distilled/deionized water (*41*).

Germ-free larvae

The $w^{1118}$ flies were obtained from the Bloomington Stock Center (Bloomington, IN) and used as the standard strain. Adult flies were maintained in autoclaved standard yeast medium containing tetracycline (50 µg/ml) for three generations to eliminate the common intracellular bacteria *Wolbachia pipientis*. Embryos of the flies were then bleached using 2.7% sodium hypochlorite and washed twice in 70% ethanol, followed by three washes in sterile water.

Washed embryos were transferred to autoclaved yeast vials without tetracycline. The third instar larvae were collected and bacterial contamination in larvae was assessed by PCR using 16S rDNA universal primers 8FE (5’-AGAGTTTGATCMTGGCTCAG-3’) and 1492R (5’-GGMTAGCTTGTTACGACTT-3’).

Cell culture

Human cervical cancer cells (HeLa) and human embryonic kidney cells (PEAKrapid) were obtained from the American Type Culture Collection (Manassas, VA). Cells were cultured in Dulbecco’s modified Eagle medium (Sigma-Aldrich, St. Louis, MO) supplemented with 10% fetal bovine serum (Thermo Fisher Scientific, Waltham, MA), 100 U/ml penicillin (Nacalai Tesque, Kyoto, Japan), and 100 µg/ml streptomycin (Nacalai Tesque) at 37°C in 5% $CO_2$.

Drosophila S2 cells (GIBCO, R690-07) were cultured at 25°C in Schneider’s medium supplemented with 10% heat-inactivated fetal bovine serum (Thermo Fisher Scientific) and 1× antibiotic-antimycotic (Thermo Fisher Scientific). Plastic Petri dishes were used for cell culture, and harvested cells were frozen at –70°C until use.

Phenotypic analysis of *Drosophila*

To analyze the effect of ppGpp on cell proliferation under starvation conditions, 50 flies that had developed for 3 days (i.e., 3 days AEL) of $w^{-}$ and *Mesh1 lof* were transferred and kept on standard or starvation medium. The number of pupae and adults in the medium were counted every 24 h.

*hs-yjbM OE* was used for the analysis of the effect of excess ppGpp. To induce *yjbM* overexpression at day 3 AEL, *hs-yjbM OE* was heat-shocked for 30 min at 37 °C and then held at 10 min at room temperature, and this heat-shock procedure was repeated. After heat shock, the larvae were cultured for 24 h on standard medium containing Formula 4-24 Instant Drosophila Medium, Blue (Carolina Biological Supply Company) at 25 °C. Larvae for the two genotypes were then gathered after incubation: *hs-FLP-GFP/w; AyGAL4, UAS-GFP/+, +/yjbM* (or +). Twenty larvae were transferred to standard medium; every 24 h, all flies were assessed as living or dead and as larvae, pupae or adults.

To observe the effects of excess ppGpp on eye development, *GMR-yjbM* adults and other control genotypes were transferred from 18 °C to 31 °C one day after emergence. After 0 (as a control) or one day incubation, the eyes and dissected eyes were examined microscopically.

### Immunohistochemistry and imaging

The experimental procedures for brain dissection, fixation, and immunostaining were carried out as described (*42*). The primary antibody for the photoreceptor cell–specific membrane protein, chaoptin, was mAb24B10 (1:50, DSHB). The secondary antibodies were conjugated to Alexa488 (1:400, Life Technologies). Images were obtained with Nikon C2+ confocal microscopes and processed with NIS-elements AR and Adobe Photoshop.

### Quantification of ppGpp and GTP by LC-ESI-MS/MS

ppGpp was extracted from cells and quantified as described by Ihara *et al*. (*23*) with slight modifications, as detailed below. Standards for ppGpp (≥85%), GTP (92%), and GTP

(>99%) were purchased from Trilink, TaKaRa, and Bio Basic Inc., respectively, and used without further purification. Fifty flies grown under normal conditions or ~65 mg of flies grown under starvation conditions were collected for all quantification except that only one larva was collected for quantification of ppGpp in *YjbM OE*. The collected samples were frozen in liquid nitrogen, broken in pieces using a pestle and mortar, and then extracted in 3 ml of 2 M formic acid/1 mM EDTA (pH 4.5). Extracts were incubated for 30 min on ice. Subsequently, 50 mM ammonium acetate/1 mM EDTA (pH 4.5) was added to the crushed tissue to a volume of 6 ml, and the extracts were divided into two 3-ml portions. To estimate the ppGpp recovery rate (*23*), 25 µl of 2 µM ppGpp was added to one of the two solutions (the standard mixed sample). For GTP quantification, this standard mixing step was omitted because the GTP recovery rate was constant perhaps because of its high concentration. To remove hydrophobic debris, 1.5 ml chloroform was added, and the mixtures were shaken three times. After centrifugation at 1600 × *g* for 1 min at room temperature, each aqueous (upper) layer was transferred to a new tube.

The samples were then subjected to solid-phase extraction. A 1-ml OASIS WAX column 30 mg (Waters) was rinsed with 1 ml ammonium acetate (pH 4.5) after 1 ml of methanol. Each sample solution was loaded onto the column, which was then washed with 1 ml of methanol after 1 ml of ammonium acetate (pH 4.5). Nucleic acids were eluted from the column with 1 ml methanol/deionized $H_2O$/25% aqueous ammonia (20:70:10) solution. The eluate was lyophilized in a Proteosave SS 15 ml Conicaltube (Sumitomo Bakelite), dissolved in 200 µl in Milli-Q water and then filtered through a FavorPrep Plasmid DNA Extraction Column (FAVORGEN). A 94-µl portion of the eluate was then mixed with 6 µl acetonitrile in a PSVial (AMR).

A 10-µl aliquot of this solution was injected into an Acquity UPLC system (Waters) and separated on an ACQUITY UPLC® BEH C18 column (2.1 × 50 mm, 1.7 µm particle size;

Waters). Nucleotides were resolved with a linear gradient of two mobile phases: solvent A (500 ml of 8 mM DMHA with 80 µl of acetic acid) and B (acetonitrile) at a flow rate of 0.3 ml $min^{-1}$, as follows: initially 0% B; 0–40% B over 10 min; 40–0% B over 10.5 min; reequilibration at 0% B over 20 min”. Effluents from UPLC were introduced on-line into an ACQUITY TQD tandem quadrupole mass spectrometer with an ESI interface (Waters). Argon was used as collision gas at 0.3 ml $min^{-1}$. Negative ESI-MS/MS detection was carried out in the multiple reaction monitoring mode or product ion scan mode using the following parameters: capillary voltage, 2.5 kV; source temperature, 150 °C; desolvation temperature, 400 °C; cone gas flow, 50 l/h; desolvation flow, 800 l/h; LM1 resolution, 13; LM 2 resolution, 15. The cone voltage and collision energy were 44 V and 44 eV, respectively, for ppGpp, and 40 V and 32 eV, respectively, for GTP. The multiple reaction monitoring transitions were 602→159 for ppGpp and 522→159 for GTP.

The ESI source was set to negative-ion mode with a capillary voltage of 2.5 kV. The source and desolvation temperatures were set to 150 and 400 °C, respectively. ppGpp was detected in multiple reaction monitoring mode. The cone voltage and collision energy were 44 V and 44 eV, respectively. The mass of the parent ion (602 *m/z*) was selected by the first quadrupole and fragmented in the collision cell to the target ion (159 *m/z*).

ppGpp and GTP levels were quantified against standard curves. Standard solutions consisted of 20 µl of 2 µM ppGpp or 0.4 mM GTP mixed with 12 µl of acetonitrile and 168 µl of Milli-Q water in a PSVial (AMR). The standard curves were linear at concentrations between 1.5625 nM and 200 nM for ppGpp, and 2500 nM and 40000 nM for GTP. The calibration curves were used when the $r^2$ values of the plotted calibration data were >0.99. The final ppGpp concentration in each sample was corrected based on its recovery rate calculated by the ppGpp standard mixed sample, as reported (*23*).

Metabolome analysis by CE-TOF MS

Ten flies of late third instar larvae were collected for metabolome analysis. Each sample was extracted in 500 µl methanol containing 8 µM of each of two reference compounds, namely methionine sulfone for the cation analysis and camphor 10-sulfonic acid for the anion analysis, using a Retsch mixer mill MM310 at a frequency of 27 Hz for 1 min. The extracts were then centrifuged at 15,000 × *g* for 3 min at 4 °C. The supernatant was transferred into a tube, and 500 µl chloroform and 200 µl water was added to the tube to perform liquid-liquid extraction. The upper layer (water) was transferred to a new tube, and evaporated for 30 min at 45°C by a centrifugal concentrator to obtain two layers. The resulting partially evaporated liquid was then centrifugally filtered through a PALL Nanosep 3-kDa cutoff filter at 9,100 × *g* for 90 min at 4°C to remove high-molecular-weight compounds such as oligo-sugars. The filtrate was evaporated to dryness for 120 min using a centrifugal concentrator. The residue (ca. 25 mg for each sample) was dissolved in 20 µl water containing 200 µM of each of the internal standards, namely 3-aminopyrrolidine for the cation analysis and trimesic acid for the anion analysis, that were used to compensate for differences in migration time in the peak annotation step.

All CE-TOF MS experiments were performed using a Model G7100A CE Instrument (Agilent Technologies, Sacramento, CA), an Agilent G6224A TOF LC/MS system, an Agilent 1200 Infinity series G1311C Quad Pump VL, an G1603A Agilent CE-MS adapter, and a G1607A Agilent CE-ESI-MS sprayer kit. The G1601BA 3D-CE ChemStation software for CE and G3335-64002 MH Workstation were used. Separations were carried out using a fused silica capillary (50 µm i.d. × 100 cm total length) filled with 1 M formic acid for cation analyses or with 20 mM ammonium formate (pH 10.0) for anion analyses as the electrolyte. The capillary temperature was maintained at 20 °C. The 15-nl sample solutions were injected at 50 mbar for 15 s. The sample tray was cooled below 10 °C. Prior to each run, the capillary was flushed with electrolyte for 5 min. The applied voltage for separation was set at 30 kV. Methanol/water (50% v/v) containing 0.5 μM reserpine was delivered as the sheath liquid at 10 μl/min. ESI-TOF MS was conducted in the positive-ion mode for cation analyses or in the negative-ion mode for anion analyses, and the capillary voltage was set at 30 kV. A flow rate of heated dry nitrogen gas (heater temperature 300 °C) was maintained at 10 l/min. The fragmentor, skimmer, and Oct RFV voltages were automatically set at optimum values. Automatic recalibration of each acquired spectrum was performed using the masses of reference standards. The methanol dimer ion ($[2M+H]^+$, *m/z* 65.0597) and reserpine ($[M+H]^+$, *m/z* 609.2806) for cation analyses or the formic acid dimer ion ($[2M-H]^-$, *m/z* 91.0037) and reserpine ($[M-H]^-$, *m/z* 607.2661) for anion analyses provided the lock mass for exact mass measurements. Exact mass data were acquired at a rate of 1.5 cycles/s over a 50–1000 *m/z* range. In every single sequence analysis (maximum 36 samples) with our CE-TOF MS system, we analyzed the standard compound mixture both before and after each analysis of a sample. The detected peak area of the standard compound mixture was compared at regular intervals to assess sensitivity and reproducibility. The standard compound mixture was composed of major detectable metabolites, including amino acids and organic acids, and this mixture was freshly prepared at least once every 6 months. For all analyses, there were no differences in the sensitivity of detection of the standard compounds mixture.

An original data file (.d) was converted to a unique binary file (.kiff) using in-house software. Peak picking and alignment were performed using another in-house software package in which peaks were picked and aligned among samples automatically. By contrast with the detected *m/z* and migration time values of standard compounds, including internal standards, peaks were annotated automatically using the same software. For normalization, the individual area of the detected peaks was divided by the peak area of the internal reference standards. Using the calibration curves for standard compounds, peak area values were converted into values corresponding to amounts. Obtained data are shown in Supplemental Dataset 1.

### Figure legends

#### Fig. S1

Comparison of MS/MS chromatograms for GTP (*m/z* = 522→159) and ppGpp (*m/z* = 602→159). GTP standards (92% or ≥99%) and larvae extracts were analyzed using the multiple reaction monitoring mode. The first quadrupole separated ppGpp (*m/z* = 602) or GTP (*m/z* = 522), and the second quadrupole separated pyrophosphate (*m/z* = 159) derived from GTP and/or ppGpp. *Multiple reaction monitoring–dependent cross talk from GTP.

#### Fig. S2.

Detection of ppGpp in various stages of *Drosophila*. Guanosine tetraphosphate–specific MS chromatograms of nucleotide pools extracted from CS eggs **(A)**, $w^{-}$ pupae **(B)**, $w^{-}$ virgin flies (male/female) **(C)**, $w^{-}$ mating flies (male/female) **(D)**, *Mesh1 lof* larvae **(E)**, and *Mesh1 lof* pupae **(F)**. The symbol + denotes the exogenous addition of ppGpp standard to the extract.

#### Fig. S3.

Concentrations of ppGpp and/or GTP in various stages of development of *Drosophila* $w^{-}$*, Mesh1 lof*, and/or *Mesh1 gof.* ppGpp levels per fresh weight (FW) **(A)**, ppGpp levels per GTP **(B)**, GTP levels per FW **(C)**, ppGpp levels per FW or GTP **(D)**, GTP levels per FW **(E)**, and GTP levels per FW in *Mesh1 gof* **(F)**. Values represent the mean ± S.D. ($n$ =3). *Significant differences ($p$ < 0.05, Student's t-test). Different letters indicate significant differences ($p$ < 0.05, Tukey's test). Male and female flies could not be distinguished at the egg and pupae stages, so they were sampled together.

**Fig. S4.**

Detection of ppGpp from various cultured cells and germ-free (GF) flies. Guanosine tetraphosphate–specific MS chromatograms of nucleotide pools extracted from flies grown under the normal or GF condition, human cultured HeLa cells, human cultured PEAKrapid cells, and *Drosophila* cultured S2 cells.

**Fig. S5.**

Starvation-induced phenotypes of *Drosophila*. Fresh weight (FW) **(A)**, ppGpp levels per GTP **(B)**, ppGpp levels per organism **(C)**, and GTP levels per FW **(D)** of $w^{-}$ and *Mesh1 lof*. Values represent the mean ± S.D. ($n$ =3). Different letters indicate significant differences ($p < 0.05$, Tukey's test). Control, flies grown under non-starvation conditions.

**Fig. S6.**

Effect of *Mesh1* mutation on eye development. Optical microscope images (left) and confocal images of dissected eyes (center and right) in $w^{-}$ **(A)** and *Mesh1 lof* **(B)**. Axons were observed by immunostaining with the chaoptin-specific antibody. Observations were made one day after emergence.

**Fig. S7.**

Effects of *Mesh1 lof* and *gof* mutations on metabolism. **(A)** Metabolites in *Mesh1 gof* larvae that were significantly changed compared with those in the control ($p < 0.05$; $n = 6$). Fold changes in metabolite levels are expressed along a color gradient. 3PG: 3-phosphoglyceric acid, 5-OxoPro: pyroglutamic acid, CML: carboxymethyl lysine, CySSG: cysteineglutathione disulfide, GAA: guanidinoacetic acid, GABA: gamma-aminobutyric acid, GSH: glutathione, GSSG: glutathione disulfide, SAM: S-adenosylmethionine, SAH: S-adenosylhomocysteine, and PEP: phosphoenolpyruvate. **(B)** Venn diagram representing the number of significantly changed metabolites in *Mesh1 lof* (left) and *Mesh1 gof* (right) compared with those of controls. Reverse means that the metabolite changes were in the opposite direction (in “reverse”; all the metabolites in *Mesh1 lof* were upregulated, and all in *Mesh1 gof* were downregulated).

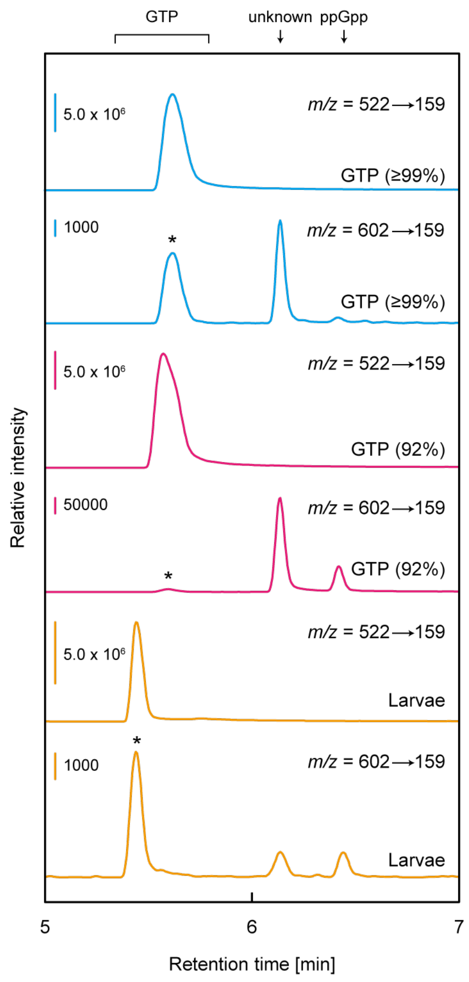
GTP
unknown
ppGpp
5.0 x 10⁶
m/z = 522→159
GTP (≥99%)
1000
m/z = 602→159
GTP (≥99%)
5.0 x 10⁶
m/z = 522→159
GTP (92%)
50000
m/z = 602→159
GTP (92%)
5.0 x 10⁶
m/z = 522→159
Larvae
1000
m/z = 602→159
Larvae
Relative intensity
5
6
7
Retention time [min]

**Fig. S1.**

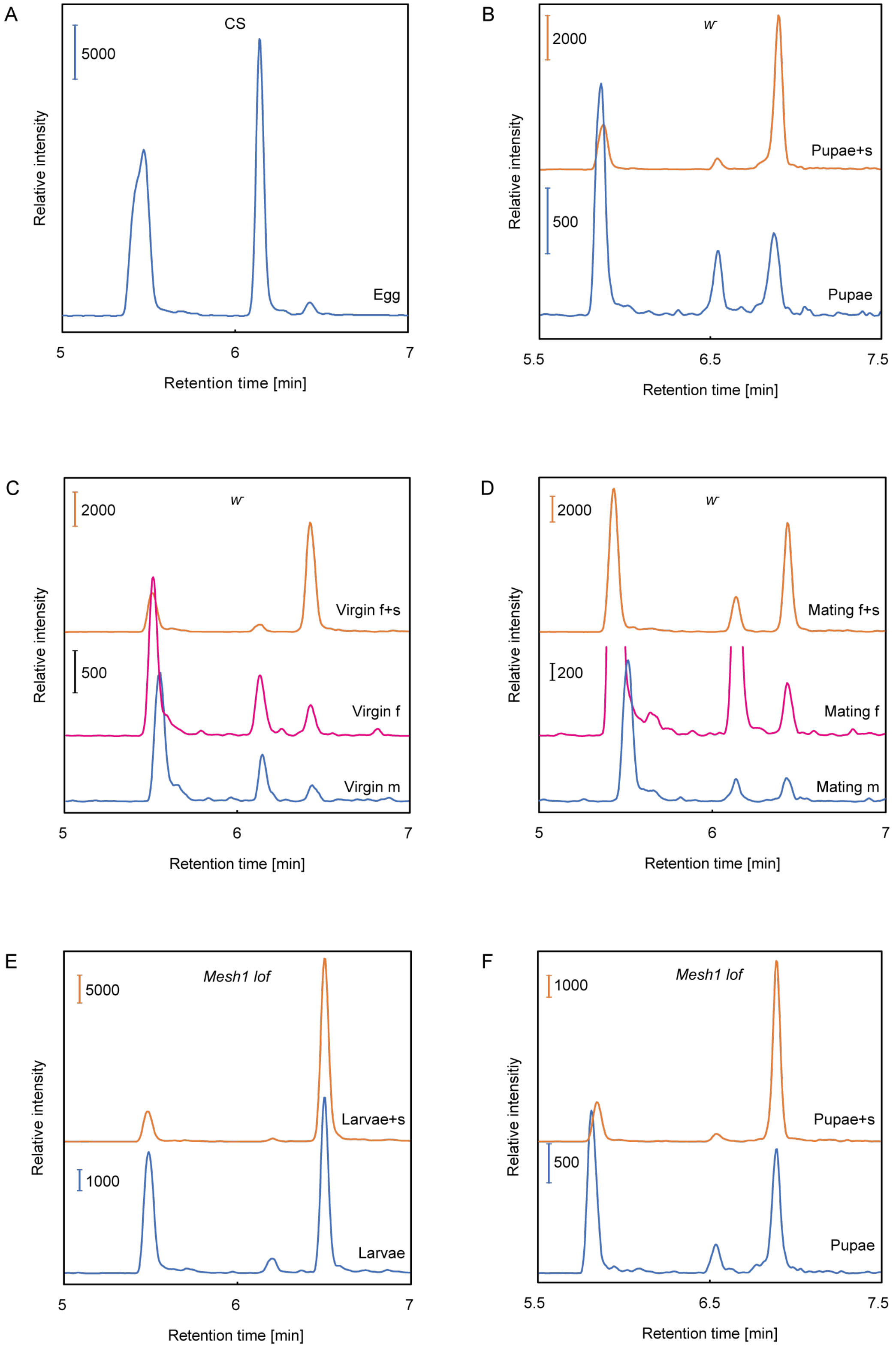
A
CS
5000
Egg
Relative intensity
5
6
7
Retention time [min]
B
w-
2000
Pupae+s
500
Pupae
Relative intensity
5.5
6.5
7.5
Retention time [min]
C
w-
2000
Virgin f+s
500
Virgin f
Virgin m
Relative intensity
5
6
7
Retention time [min]
D
w-
2000
Mating f+s
200
Mating f
Mating m
Relative intensity
5
6
7
Retention time [min]
E
Mesh1 lof
5000
Larvae+s
1000
Larvae
Relative intensity
5
6
7
Retention time [min]
F
Mesh1 lof
1000
Pupae+s
500
Pupae
Relative intensity
5.5
6.5
7.5
Retention time [min]

**Fig. S2.**

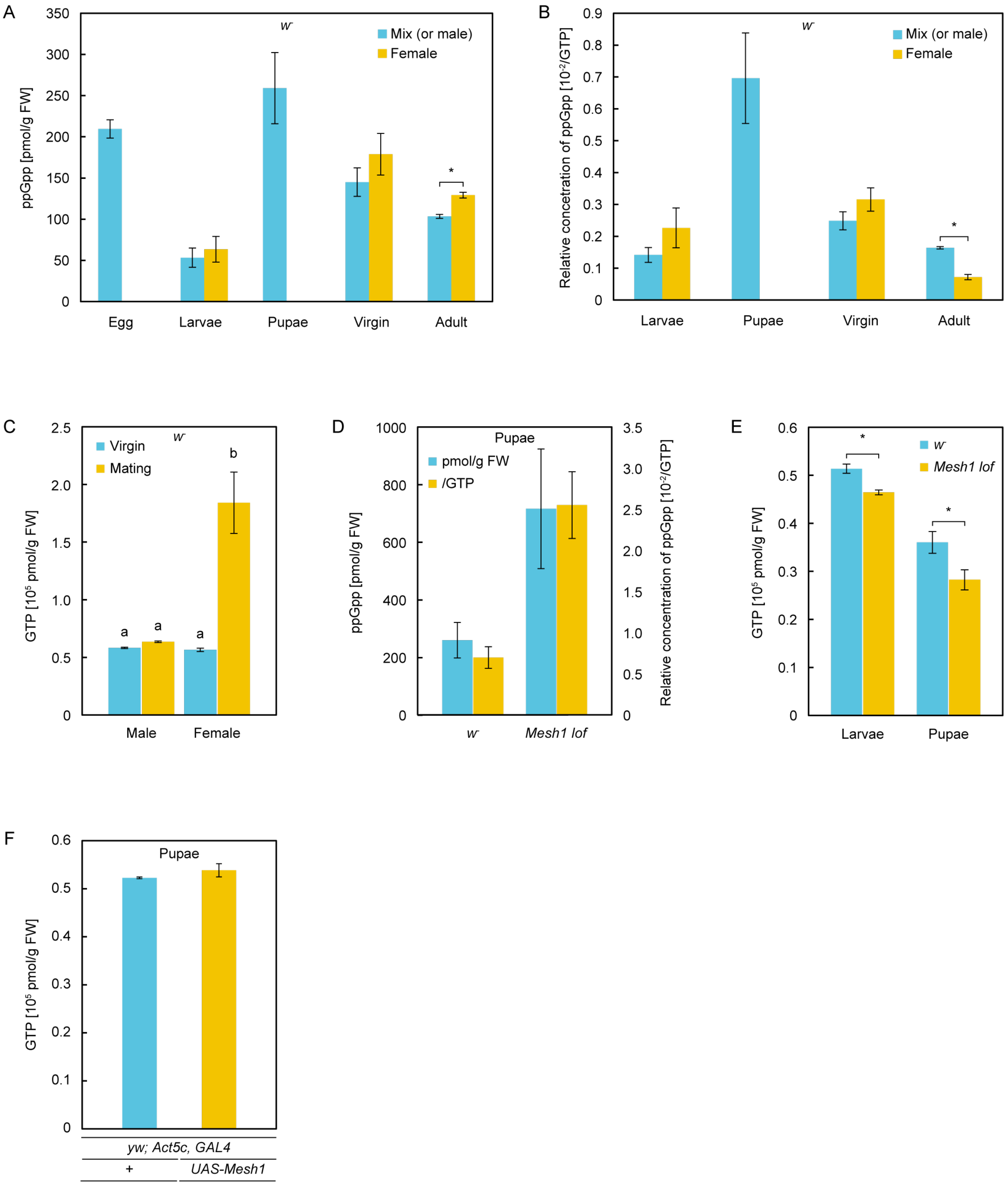
A
w-
Mix (or male)
Female
ppGpp [pmol/g FW]
Egg
Larvae
Pupae
Virgin
Adult
B
w-
Relative concetration of ppGpp [10-2/GTP]
Larvae
Pupae
Virgin
Adult
C
w-
Virgin
Mating
GTP [10^5 pmol/g FW]
Male
Female
D
Pupae
pmol/g FW
/GTP
ppGpp [pmol/g FW]
Relative concentration of ppGpp [10-2/GTP]
w-
Mesh1 lof
E
w-
Mesh1 lof
GTP [10^5 pmol/g FW]
Larvae
Pupae
F
Pupae
GTP [10^5 pmol/g FW]
yw; Act5c, GAL4
+
UAS-Mesh1

**Fig. S3.**

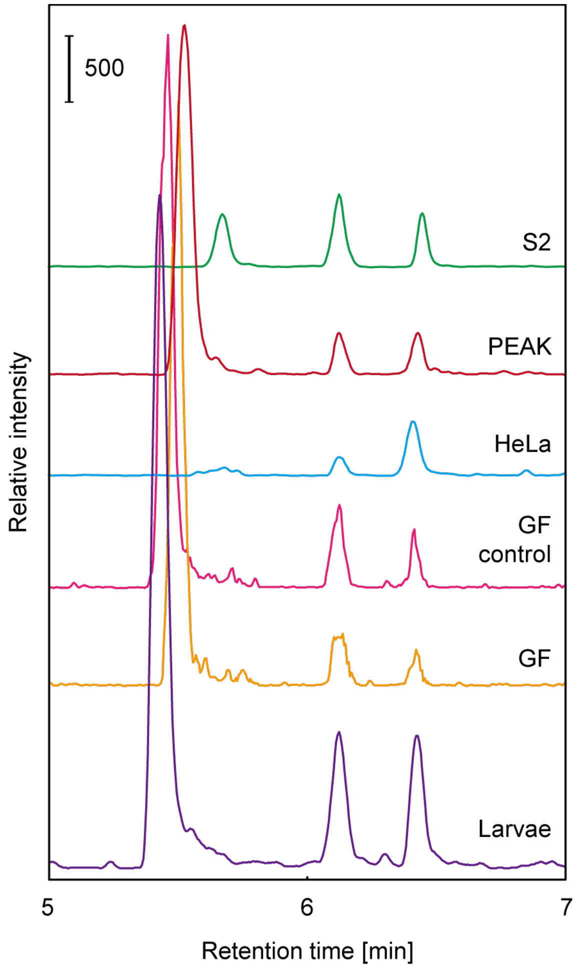
500
S2
PEAK
HeLa
GF
control
GF
Larvae
Relative intensity
5
6
7
Retention time [min]

**Fig. S4.**

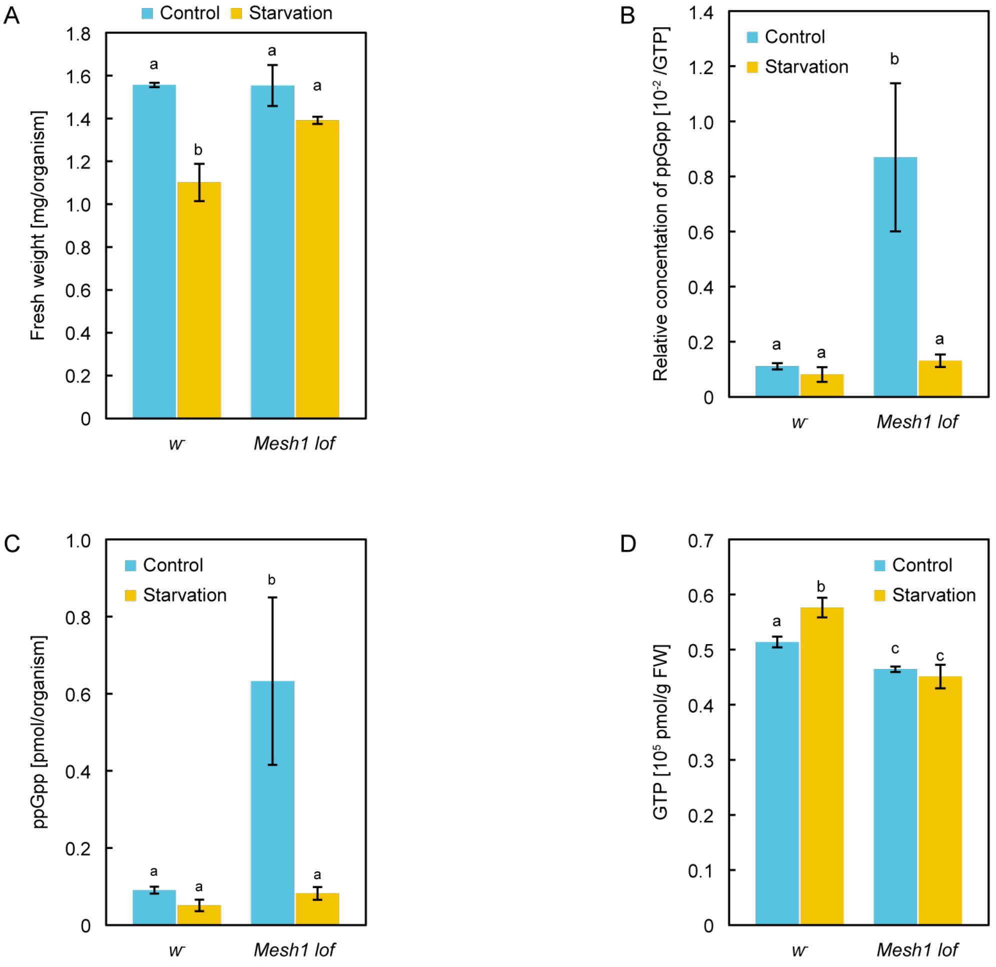
A
Control
Starvation
Fresh weight [mg/organism]
1.8
1.6
1.4
1.2
1.0
0.8
0.6
0.4
0.2
0
a
b
a
a
w-
Mesh1 lof
B
Control
Starvation
Relative concentation of ppGpp [10-2 /GTP]
1.4
1.2
1.0
0.8
0.6
0.4
0.2
0
a
a
b
a
w-
Mesh1 lof
C
Control
Starvation
ppGpp [pmol/organism]
1.0
0.8
0.6
0.4
0.2
0
a
a
b
a
w-
Mesh1 lof
D
Control
Starvation
GTP [105 pmol/g FW]
0.7
0.6
0.5
0.4
0.3
0.2
0.1
0
a
b
c
c
w-
Mesh1 lof

**Fig. S5.**

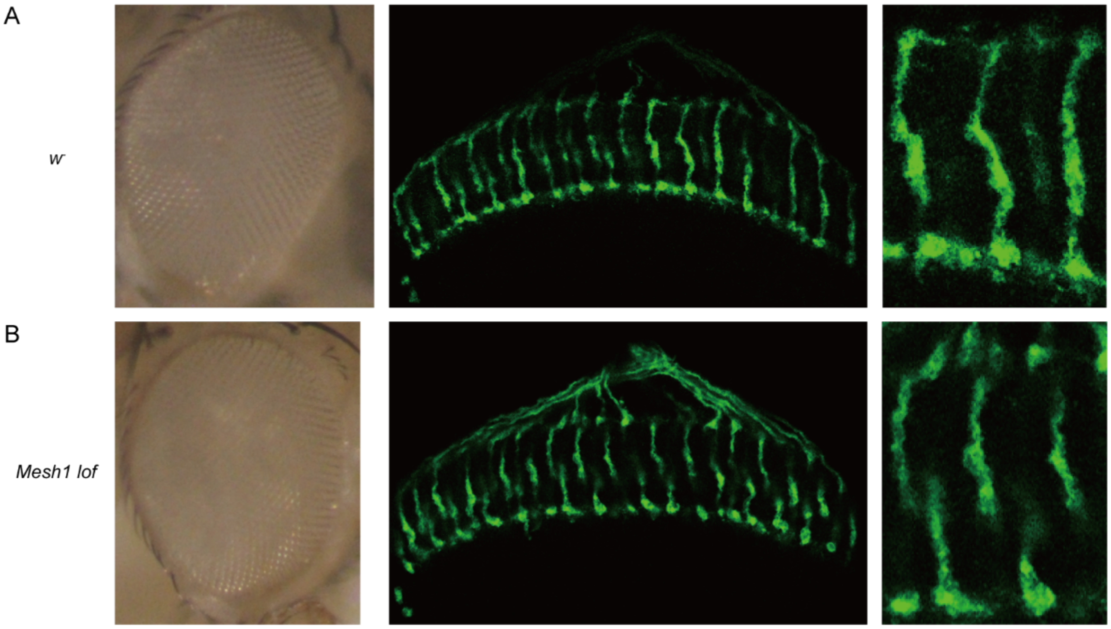

**Fig. S6.**

A *Mesh1 gof*/con.

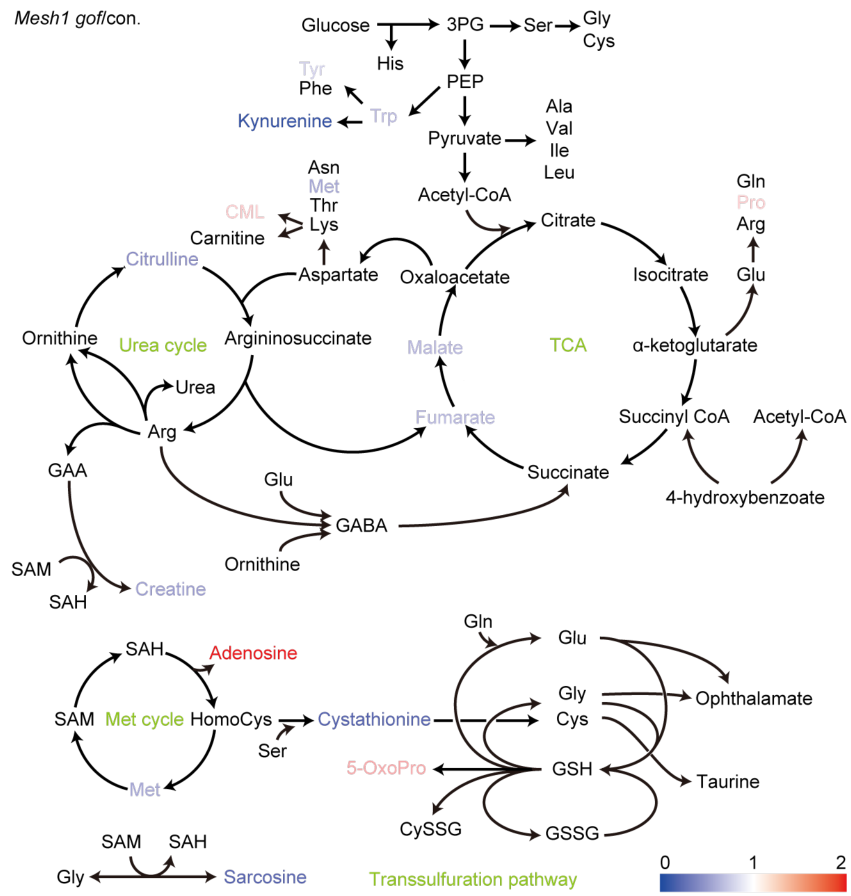

B

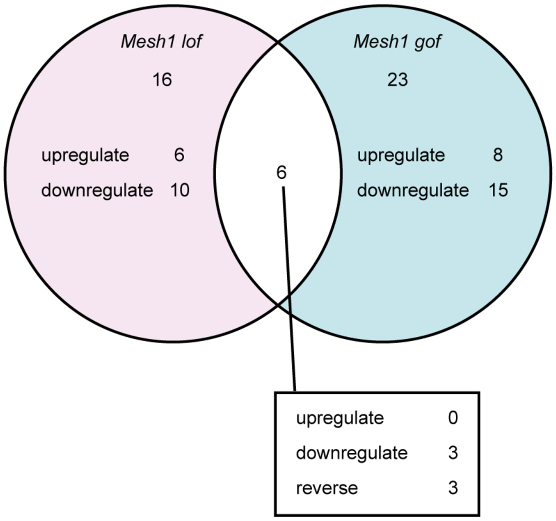

**Fig. S7.**

**Table S1.**

ppGpp levels (per FW or GTP) in each stage. Values represent the mean ± S.D. ($n$ =3). N.D., not determined.

| Species | pmol/g FW | S.D. | ($10^{-3}$)/GTP | S.D. ($10^{-3}$) |
|---|---|---|---|---|
| ***Drosophila melanogaster*** | | | | |
| **CS (WT)** | | | | |
| Eggs | 209.6 | 11.1 | N.D. | N.D. |
| ***w*⁻ (WT)** | | | | |
| Third instar larvae (male) | 53.3 | 11.7 | 1.41 | 0.231 |
| Third instar larvae (female) | 63.5 | 15.6 | 2.27 | 0.624 |
| Starved larvae | 47.1 | 16.9 | 0.812 | 0.266 |
| Pupae (day 2) | 259 | 43.2 | 6.96 | 1.42 |
| Virgin (male) | 145 | 17.3 | 2.49 | 0.282 |
| Virgin (female) | 179 | 25.2 | 3.16 | 0.367 |
| Mating adults (male, day 4) | 103 | 2.44 | 1.63 | 0.0372 |
| Mating adults (female, day 4) | 129 | 3.49 | 0.710 | 0.0830 |
| ***Mesh1 lof*** | | | | |
| Third instar larvae | 403 | 121 | 8.61 | 2.64 |
| Starved larvae | 59.1 | 11.3 | 1.31 | 0.23 |
| Pupae (day 2) | 716 | 223 | 25.5 | 8.31 |
| *yjbM OE* (day 3 larvae) | 45969 | 12183 | 185.3 | 96.6 |
| *Mesh1 gof* (day 2 pupae) | 347 | 112 | 6.5 | 2.2 |
| Germ-free larvae | 24.8 | 5.9 | 0.129 | 0.0155 |
| **Cultured cells** | | | | |
| HeLa | 43.5 | 12.6 | 2.54 | 0.253 |
| S2 | N.D. | N.D. | 2.74 | 0.780 |

**Table S2.**

Metabolites that were present in significantly different amounts in *Mesh1 lof* compared with WT. Third instar larvae grown in standard medium were used for metabolite analyses. For each metabolite, the relative increase or decrease and *p* value are indicated ($n = 6$). GABA: gamma-aminobutyric acid, CysSG: cysteineglutathione disulfide, HMG: β-hydroxy β-methylglutaryl-CoA, GlcNAc1P: N-acetyl-α-D-glucosamine 1-phosphate, NADP: nicotinamide adenine dinucleotide phosphate.

| Annotation Name | *Mesh1 lof*/WT | *p* value |
|---|---|---|
| 4-Hydroxybenzoate | 1.87 | $1.75 \times 10^{-2}$ |
| Fumarate | 1.45 | $5.00 \times 10^{-2}$ |
| GABA | 1.43 | $3.13 \times 10^{-2}$ |
| Malate | 1.40 | $2.98 \times 10^{-3}$ |
| His | 1.35 | $1.58 \times 10^{-2}$ |
| CysSG | 1.34 | $4.97 \times 10^{-2}$ |
| Raffinose | 1.31 | $2.47 \times 10^{-2}$ |
| Methylneuraminate | 1.27 | $2.69 \times 10^{-2}$ |
| Carnitine | 1.23 | $1.20 \times 10^{-3}$ |
| HMG | 0.77 | $9.56 \times 10^{-3}$ |
| Arg | 0.76 | $2.18 \times 10^{-2}$ |
| Gly-Leu | 0.64 | $1.50 \times 10^{-2}$ |
| GlcNAc1P | 0.64 | $8.40 \times 10^{-4}$ |
| Lys | 0.60 | $3.50 \times 10^{-2}$ |
| Taurine | 0.55 | $1.88 \times 10^{-5}$ |
| NADP | 0.50 | $4.96 \times 10^{-2}$ |
| Cytosine | 0.48 | $7.17 \times 10^{-3}$ |
| Sedoheptulose 7-phosphate | 0.44 | $3.23 \times 10^{-3}$ |
| Deoxyguanosine | 0.39 | $4.13 \times 10^{-2}$ |
| Sarcosine | 0.31 | $2.96 \times 10^{-4}$ |
| Kynurenine | 0.22 | $9.15 \times 10^{-3}$ |
| 1-Methyladenosine | 0.12 | $9.53 \times 10^{-5}$ |

**Table S3.**

Metabolites that were present in significantly different amounts in *Mesh1 gof* compared with WT. Third instar larvae grown in standard medium were used for metabolite analyses. For each metabolite, the relative increase or decrease and *p* value are indicated (*n* = 6). CDP: cytidine diphosphate, 5-OxoPro: pyroglutamic acid, PhenylP: phenylephrine, CML carboxymethyl lysine, 3-MethylHis: 3-methylhistidine, HomoPro: homoproline, GlcNAc1P: N-acetyl-α-D-glucosamine 1-phosphate, 2-MethylSer: 2-methylserine, MethylAla: methylalanine.

| Annotation Name | *Mesh1 gof* /con. | *p* value |
|---|---|---|
| Adenosine | 2.10 | $3.37 \times 10^{-2}$ |
| Adenine | 1.62 | $1.87 \times 10^{-2}$ |
| CDP | 1.62 | $2.25 \times 10^{-2}$ |
| 5-OxoPro | 1.32 | $2.14 \times 10^{-2}$ |
| Pyridoxamine5P | 1.26 | $2.81 \times 10^{-2}$ |
| CML | 1.19 | $2.81 \times 10^{-3}$ |
| PhenylP | 1.17 | $1.66 \times 10^{-2}$ |
| Pro | 1.15 | $2.01 \times 10^{-2}$ |
| 3-MethylHis | 0.86 | $4.23 \times 10^{-2}$ |
| Tyr | 0.84 | $6.30 \times 10^{-4}$ |
| Deoxyuridine | 0.79 | $4.72 \times 10^{-4}$ |
| HomoPro | 0.75 | $1.87 \times 10^{-2}$ |
| Trp | 0.74 | $2.03 \times 10^{-2}$ |
| Malate | 0.74 | $1.05 \times 10^{-2}$ |
| Methylneuraminate | 0.73 | $9.58 \times 10^{-3}$ |
| Fumarate | 0.73 | $1.02 \times 10^{-2}$ |
| Fructose-Pro | 0.72 | $5.95 \times 10^{-4}$ |
| Fructose-Ala | 0.71 | $1.34 \times 10^{-3}$ |
| GlcNAc1P | 0.71 | $3.63 \times 10^{-4}$ |
| N8-Acetylspermidine | 0.71 | $2.52 \times 10^{-2}$ |
| Met | 0.68 | $6.90 \times 10^{-3}$ |
| Hypoxanthine | 0.67 | $5.45 \times 10^{-3}$ |
| Creatine | 0.64 | $9.51 \times 10^{-4}$ |
| Citrulline | 0.57 | $3.01 \times 10^{-3}$ |

| | | |
|---|---|---|
| 2-MethylSer | 0.55 | $2.42 \times 10^{-2}$ |
| Sarcosine | 0.42 | $2.21 \times 10^{-5}$ |
| Cystathionine | 0.41 | $3.93 \times 10^{-3}$ |
| MethylAla | 0.25 | $2.12 \times 10^{-5}$ |
| Kynurenine | 0.23 | $3.83 \times 10^{-3}$ |
